# Elevated expression of *ACE2* in tumor-adjacent normal tissues of cancer patients

**DOI:** 10.1101/2020.04.25.061200

**Authors:** Tom Winkler, Uri Ben-David

**Affiliations:** Department of Human Molecular Genetics and Biochemistry, School of Medicine, Tel Aviv University, Tel Aviv, Israel

## Abstract

The rapidly developing COVID-19 pandemic has raised a concern that cancer patients may have increased susceptibility to SARS-CoV-2 infection. This discussion has mostly focused on therapy-induced immune suppression. Here, we examined the expression patterns of *ACE2*, the receptor through which SARX-CoV2 enters human cells, and found that *ACE2* mRNA levels are elevated in tumor-adjacent normal tissues of cancer patients, including in normal-adjacent lung tissues of lung cancer patients. These observations raise the possibility that the elevated COVID-19 risk of cancer patients may not be limited to those undergoing immune-suppressing treatment.

---

The recent outbreak of a novel betacoronavirus known as severe acute respiratory syndrome corona virus 2 (SARS-CoV-2) has raised the concern that cancer patients might be particularly susceptible to infection by this virus^1,2,3^. Importantly, the guidelines for cancer patients during the COVID-19 pandemic focus on lung cancer patients who are undergoing active chemotherapy or radical radiotherapy, and on patients with blood cancers^1^. Intentional postponing of adjuvant chemotherapy or elective surgery for stable cancer has even been proposed to alleviate the risk^2^. However, whether patients with other epithelial solid tumors are also more susceptible to COVID-19 is currently unknown.

SARS-CoV-2 requires the Angiotensin Converting Enzyme 2 (ACE2) to enter human cells^4,5^. Moreover, *ACE2* gene expression levels in epithelial tissues corresponded to survival following SARS-CoV infection in transgenic mice^6^, and soluble human ACE2 inhibited SARS-CoV-2 infections in engineered human tissues^7^. *ACE2* mRNA levels are particularly high in the human kidney, testis, heart and intestinal tract^8,9^. Albeit not highly expressed in the normal lungs, *ACE2* expression levels in the airway epithelia increase due to chronic exposure to cigarette smoke^8,10^, which was associated with infection susceptibility^11^. *ACE2* expression levels have also been suggested to underlie the increased susceptibility of patients with hypertension and diabetes to SARS-CoV2 infection^12^, and to be increased in patients with co-morbidities associated with severe COVID-19^13^. Therefore, *ACE2* expression in epithelial tissues, and in particular in the epithelial airway, seem to have considerable effect on COVID-19 morbidity and mortality.

We compared *ACE2* mRNA levels between normal tissues (NT), primary tumors (PT), and normal tissues adjacent to tumors (NAT), using data from The Cancer Genome Atlas (TCGA) and GTEx^14^. In almost all of the relevant tissues for which data were available, *ACE2* mRNA levels in PT were significantly higher than in NT of the respective tissue (Figure 1 and Supplementary Figure 1). Surprisingly, *ACE2* expression levels in NAT were also significantly higher than in NT across tissues, and in all cases were at least as high as in the respective PT (Figure 1 and Supplementary Figure 1). This result suggests that the NAT of cancer patients would likely be more susceptible to SARS-CoV2 infection than the corresponding tissues of healthy individuals.

**Figure 1:**
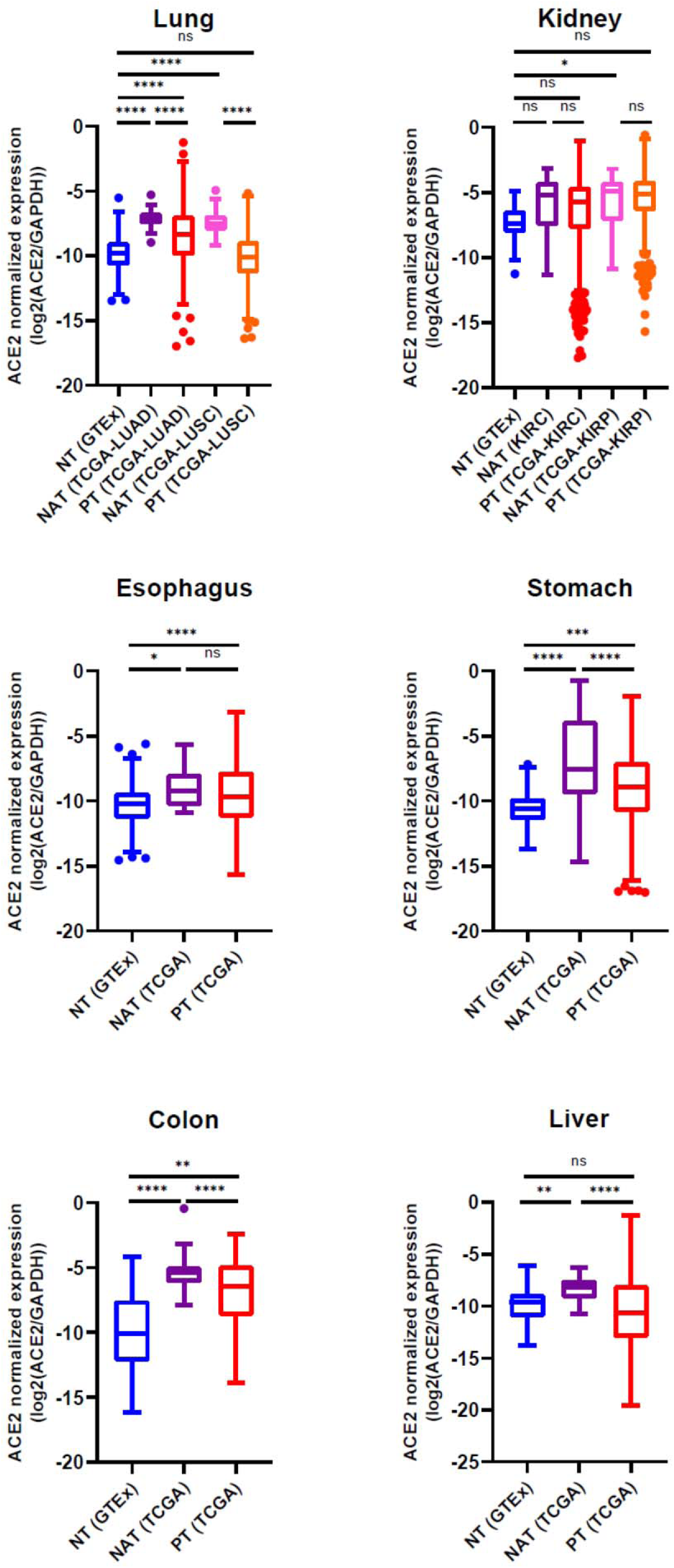
Elevated expression of *ACE2* in tumor-adjacent normal tissues of cancer patients. mRNA levels of *ACE2* (normalized to GAPDH) in normal tissues (NT), Normal adjacent tissues (NAT) and primary tumors (PT) across 8 tumor types representing 6 distinct tissues. LUAD, lung adenocarcinoma; LUSC, lung squamous cell carcinoma; KIRC, Kidney renal clear cell carcinoma; KIRP, kidney renal papillary cell carcinoma. Bar, median; box, 25th and 75th percentiles; whiskers, 1.5 × IQR of lower and upper quartile. *, p<0.05; **, p<0.01; ***, p<0.001; ****, p<0.0001; One-way ANOVA with Tukey’s test.

Focusing on the lung due to its relevance in the disease etiology, we next queried the mRNA expression levels of *ACE2* in two additional datasets of normal human tissues, the Human Protein Atlas^15^ and FANTOM5^16^. In concordance with the GTEx data, the expression levels of ACE2 in whole-lung tissues from healthy donors were negligible (median of 0.7pTPM, 1.8pTPM and 2.6 scaled tags per million, in GTEx, HPA and FANTOM5, respectively). Next, we compared the relative expression levels of *ACE2* between healthy and tumor-adjacent lung tissues, using 6 published gene expression microarray datasets^17–20^. ACE2 expression levels in the tumor-adjacent normal lung samples were detected at discernible levels, and were significantly higher than those in the healthy normal lung samples (Figure 2). This analysis confirmed that the mRNA levels of *ACE2* are elevated in tumor-adjacent lungs of lung cancer patients.

**Figure 2:**
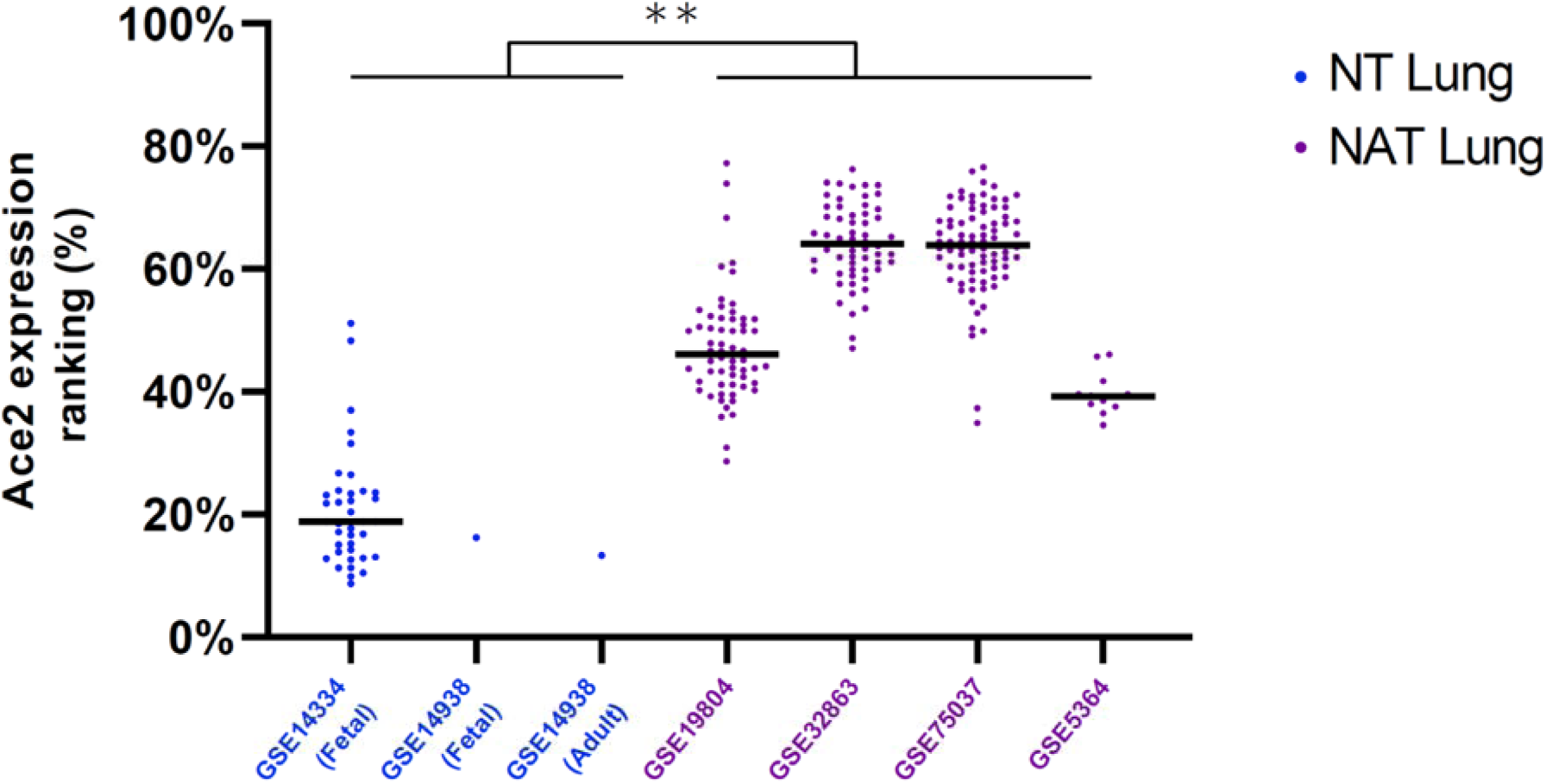
Elevated expression of ACE2 in multiple studies of tumor-adjacent lung tissues from lung cancer patients. Relative mRNA levels of *ACE2* in normal lung tissues (NT) and normal adjacent tissues (NAT), calculated based on 6 gene expression microarray datasets. Bar, median. **, p=0.0045; two-tailed Student’s t-test for a comparison between the median values of the datasets in each group.

This observation raises the possibility that lung cancer patients may have an increased risk to SARS-CoV2 infection, regardless of chemotherapy-induced immune suppression. Furthermore, patients with other types of cancer, such as renal or gastrointestinal cancers, may also have elevated infection risk. However, to determine whether this is indeed the case, two questions require urgent attention: 1) Is *ACE2* expression level in non-lung epithelia associated with SARS-coV2 infection risk?; and 2) Is *ACE2* upregulation limited to the tissue adjacent to the tumor (presumably due to the tumor microenvironment), or are *ACE2* levels systemically elevated in cancer patients? Until these questions are resolved, we propose that the discussion of cancer guidelines during the COVID-19 pandemic should expand beyond patients with treatment-induced immune suppression.

## Methods

GTEx and TCGA expression data were extracted from UCSC Xena^14^ (https://xena.ucsc.edu/). For each tumor, the expression of each gene was defined as log2(expected_count-deseq2+1). The normalized expression of ACE2 is either presented as is (Supplementary Fig. 1) or further normalized by subtracting the normalized expression of GAPDH from the normalized expression of ACE2 (Fig. 1).

mRNA expression data of lung tissues were obtained from the Gene Expression Omnibus (https://www.ncbi.nlm.nih.gov/geo/) using the following accession numbers: GSE14334, GSE14938, GSE5364, GSE19804, GSE32863 and GSE75037. ACE2 probeset ID’s were obtained from the annotation files of the respective microarray platforms. For each dataset, the expression levels of all probesets were ranked within each sample, and the rank of the highest-expressing *ACE2* probeset was considered.

### Statistical analysis

Statistical analyses were performed using PRISM GraphPad 8.4.1. One-way ANOVA with Tukey’s multiple comparison test was used to determine the statistical significance of the differences in gene expression between the groups. Two-tailed Student’s t-test on the median ACE2 expression percentile values was used to determine the statistical significance of the ranking difference between the groups.

---

**Supplementary Figure 1:**
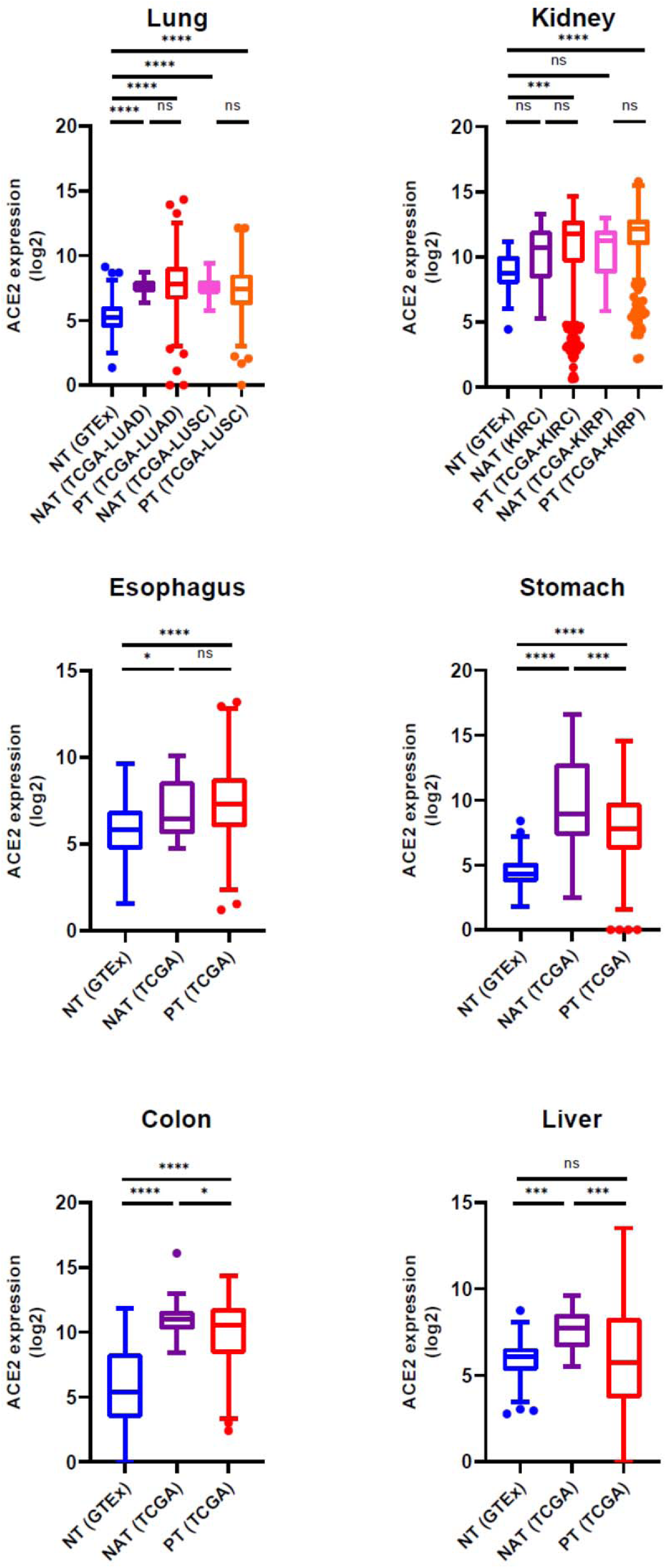
Elevated expression of *ACE2* in tumor-adjacent normal tissues of cancer patients. mRNA levels of ACE2 (without normalization to GAPDH) in normal tissues (NT), Normal adjacent tissues (NAT) and primary tumors (PT) across 8 tumor types representing 6 distinct tissues. LUAD, lung adenocarcinoma; LUSC, lung squamous cell carcinoma; KIRC, Kidney renal clear cell carcinoma; KIRP, kidney renal papillary cell carcinoma. Bar, median; box, 25th and 75th percentiles; whiskers, 1.5 × IQR of lower and upper quartile. *, p<0.05; **, p<0.01; ***, p<0.001; ****, p<0.0001; One-way ANOVA with

